# Engineered RecA constructs reveal the minimal SOS activation complex

**DOI:** 10.1101/2022.09.30.510415

**Authors:** Michael B. Cory, Allen Li, Christina M. Hurley, Zachary M. Hostetler, Yarra Venkatesh, Chloe M. Jones, E. James Petersson, Rahul M. Kohli

**Affiliations:** Graduate Group in Biochemistry and Biophysics, University of Pennsylvania, Philadelphia, Pennsylvania 19104, United States; Department of Medicine, University of Pennsylvania, Philadelphia, Pennsylvania 19104, United States; Department of Chemistry, University of Pennsylvania, Philadelphia, Pennsylvania 19104, United States; Department of Biochemistry and Biophysics, University of Pennsylvania, Philadelphia, Pennsylvania 19104, United States

**Keywords:** RecA, LexA, SOS, Synthetic Biology, Protein Engineering, Antibiotic Resistance

## Abstract

The SOS response is a bacterial DNA damage response pathway that has been heavily implicated in bacteria’s ability to evolve resistance to antibiotics. Activation of the SOS response is dependent on the interaction between two bacterial proteins, RecA and LexA. RecA acts as a DNA damage sensor by forming lengthy oligomeric filaments (RecA*) along single-stranded DNA (ssDNA) in an ATP-dependent manner. RecA* can then bind to LexA, the repressor of SOS response genes, triggering LexA degradation and leading to induction of the SOS response. Formation of the RecA*-LexA complex therefore serves as the key ‘SOS activation signal’. Given the challenges associated with studying a complex involving multiple macromolecular interactions, the essential constituents of RecA* that permit LexA cleavage are not well defined. Here, we leverage head-to-tail linked and end-capped RecA constructs as tools to define the minimal RecA* filament that can engage LexA. In contrast to previously postulated models, we found that as few as three linked RecA units are capable of ssDNA binding, LexA binding, and LexA cleavage. We further demonstrate that RecA oligomerization alone is insufficient for LexA cleavage, with an obligate requirement for ATP and ssDNA binding to form a competent SOS activation signal with the linked constructs. Our minimal system for RecA* highlights the limitations of prior models for the SOS activation signal and offers a novel tool that can inform efforts to slow acquired antibiotic resistance by targeting the SOS response.

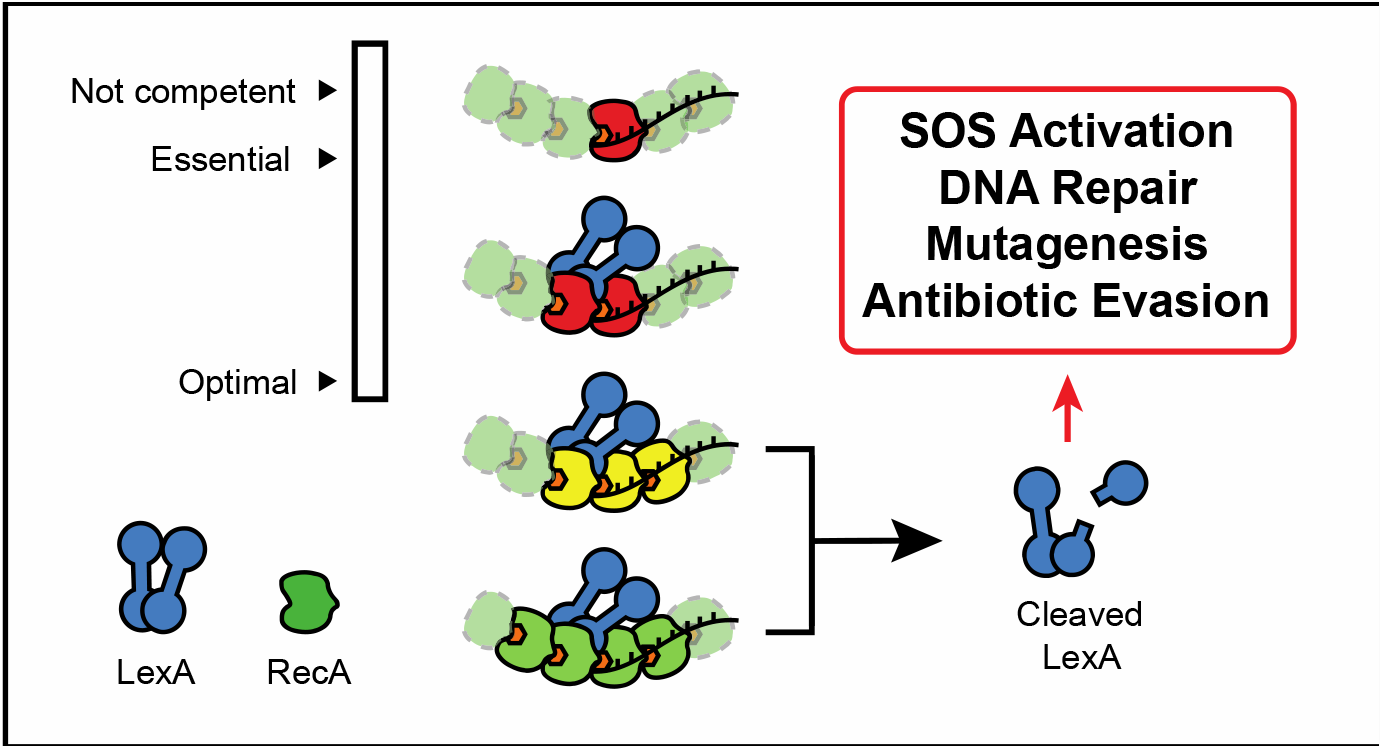

## INTRODUCTION

Across most bacterial species, a tightly regulated stress response pathway known as the SOS response mediates a robust ability to adapt to environmental stressors that threaten genomic integrity.^1,2^ In *E. coli*, the SOS response involves over 50 identified genes that serve to sense and respond to DNA damage.^3-6^ Pathway activation includes early induction of various DNA repair enzymes, allowing for high-fidelity removal and repair of DNA lesions.^6,7^ Prolonged SOS activation, in contrast, promotes a transient hypermutator state that has been linked to the SOS-driven overexpression of pro-mutagenic translesion polymerases PolIV and PolV.^8-10^ Given its physiological roles, the SOS response can also impact multiple mechanisms relevant to antibiotic therapy. Genetic inactivation of the SOS response results in hypersensitization of bacteria to antibiotics that induce DNA damage.^11-14^ Additionally, SOS deficiency greatly diminishes the rate of acquired resistance via hypermutation.^11,12^ Characterization of the molecular events leading to SOS activation can therefore offer insights into bacterial adaptation and evolution, which may consequently inform new potential strategies for combating antibiotic resistant bacteria.

The ubiquity of threats to genomic integrity explains the highly conserved nature of the two key regulatory proteins – RecA and LexA – whose interaction governs SOS activation (**Fig. 1**). LexA is a homodimeric repressor that binds operator sites within the promoters of SOS genes in the absence of DNA damage.^15-19^ RecA, named for its role in homologous recombination, acts as a sensor for single-stranded DNA (ssDNA) that accumulates during stalled replication or through processing of single- or double-stranded DNA breaks associated with DNA damage.^20-26^ RecA binds to ssDNA in an ATP-dependent manner, forming a helical nucleoprotein polymer termed RecA*, which is considered the biologically active species involved in homologous recombination and SOS response activation.^27-32^ RecA* can bind to free LexA dimers, forming the SOS activation complex and triggering an auto-proteolytic cleavage event in LexA that severs its DNA binding N-terminal domain (NTD) from its dimerization and catalytic core C-terminal domain (CTD).^33-36^ The depletion of free LexA leads to the ordered de-repression of SOS pathway genes as DNA-bound LexA dissociates, and a cascade of DNA repair and DNA damage tolerance activities are induced.^5,6^ As DNA damage is reversed, RecA* is depleted and LexA, which is autoregulated, reaccumulates to restore SOS pathway repression.^15,37^

**Figure 1.**
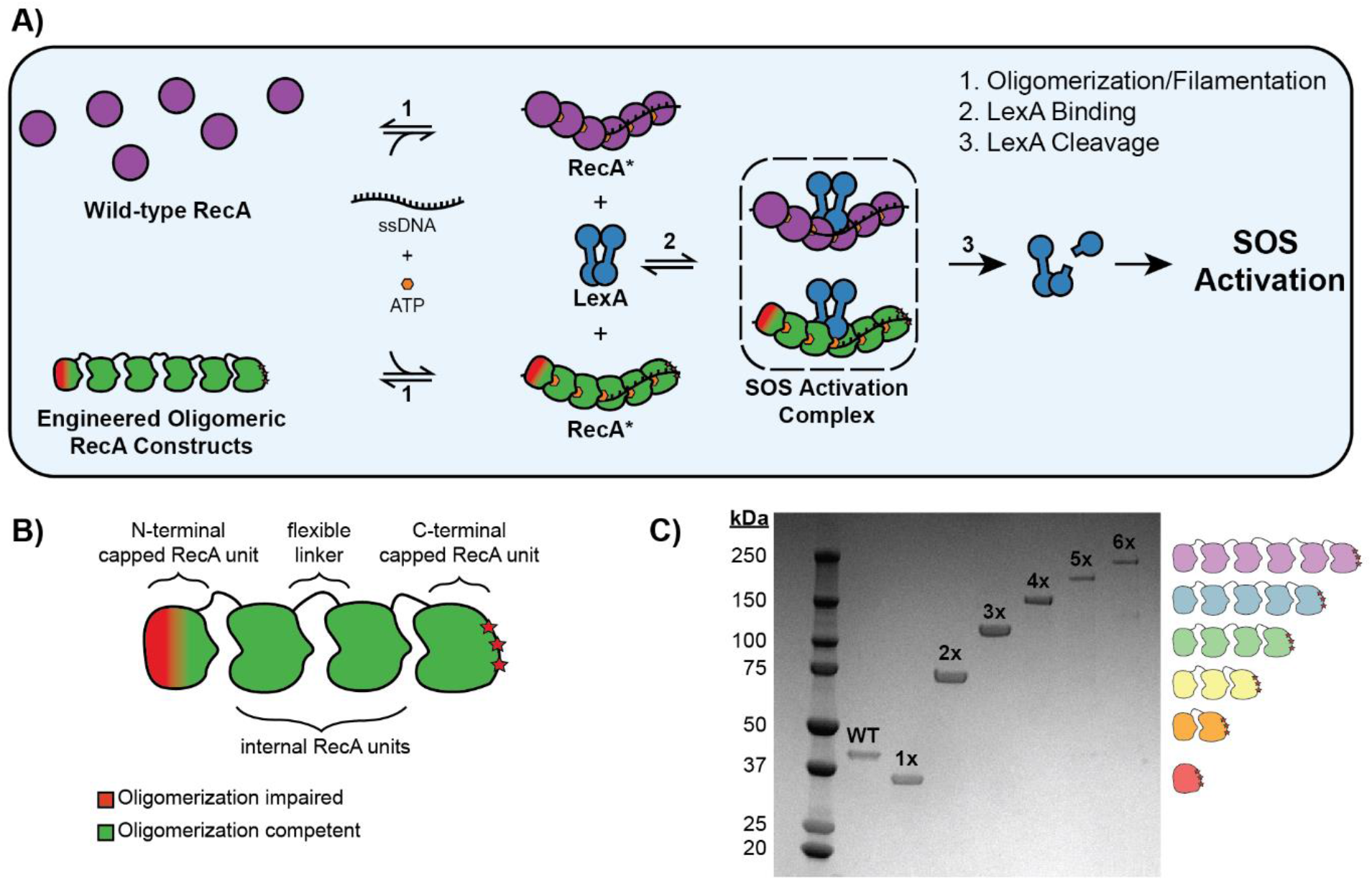
Design of engineered RecA oligomeric constructs used to probe SOS pathway activation. **A)** Reaction schema of three major steps in SOS pathway activation: (1) RecA* formation marked by RecA filamentation, (2) LexA binding to form the RecA*-LexA or SOS activation complex, and (3) LexA cleavage. **B)** Graphical representation of engineered RecA constructs. Alterations that impair further oligomerization from the N- and C-terminal ends are colored red. **C)** 4-20% SDS-PAGE gel of each construct after expression and purification.

The assembly of large macromolecular complexes is inherently challenging to study, and the molecular details regarding the formation of the SOS activation complex have therefore remained elusive.^38-44^ These challenges are compounded by several factors specific to the SOS pathway. For one, RecA has at least two distinct functional roles – catalyzing homologous recombination and serving as a co-protease for LexA – making the interpretation of mutational studies difficult.^45^ Second, RecA can also form a variety of heterogeneous macromolecular structures in solution, ranging from long and ordered RecA* filaments to non-filamentous oligomeric forms lacking ATP or DNA.^46,47^ Given the challenges posed by RecA, several proposed models of RecA* binding to LexA have emerged with some shared features and others that conflict. While previous models consistently suggest that LexA binding takes place within the groove of the RecA* filament, major points of disparity remain relating to RecA:LexA stoichiometry and the identification of specific residues that may be important for binding.^48-50^ Early transmission electron microscopy structural models predicted that between two and three RecA units can bind to LexA, whereas evolutionary trace mutagenesis and chemical crosslinking results suggested that as many as seven consecutive RecA protomers are needed to form the interaction interface.^48,50^ Implicit in these models is the assumption that the active RecA* filament is the only form of RecA capable of binding to LexA, despite the existence of non-ssDNA-bound RecA oligomers that can form both *in vitro* and *in vivo*.^46,51^ The uncertain nature of the size of the LexA binding interface on RecA* limits efforts to robustly define and evaluate a kinetic model of SOS activation. Developing a simpler and more mutable system could help overcome the above challenges that preclude elucidation of the molecular basis for SOS activation.

Here, we employ engineered RecA constructs as a simplified system to define the essential constituents for the RecA*-LexA complex. Building on precedents whereby constructs of defined oligomer size have been used to solve the structure of RecA*,^52^ we generate and characterize a series of engineered RecA oligomers ranging from one to six translationally-linked protomers (RecA1x – 6x). This approach allows us to systematically probe the biochemical requirements of LexA cleavage using fluorescence-based approaches that inform on each individual step in the SOS activation process (**Fig. 1**). Our work here both enables us to define the essential requirements for a RecA* species to support LexA cleavage and also offers a novel tool for further biochemical and structural exploration of SOS activation.

## MATERIALS AND METHODS

### Design of Engineered Oligomeric RecA Constructs

The sequence design for each *recA* protomer gene was based on the wild-type *recA* sequence from *E. coli* strain K12. To facilitate site-directed mutagenesis, allow for protomer-specific sequencing, and minimize *in vivo* recombination of the *recA* protomers during plasmid assembly, the sequence for each protomer was diversified using an online codon optimization tool (Integrated DNA Technologies, IDT). Each *recA* protomer sequence contained modifications that were previously demonstrated to support the assembly of length-restricted nucleoprotein filaments.^52,53^ These modifications include the removal of the C-terminal amino acids (residues 336-353) in all *recA* protomers, the removal of the α-β oligomerization motif (residues 1-30) from the first (N-terminal) *recA* protomer, and the substitution of three amino acids (C117M, S118V, and Q119R) in the last (C-terminal) *recA* protomer. A polyhistidine tag was included on the N-terminal *recA* protomer and the protomers were linked with a fourteen amino acid flexible linker (TGSTGSGTTGSTGS). Finally, unique restriction enzyme sites were introduced to allow for facile ligation of the gene fragments in downstream sub-cloning steps (**Fig. S1**).

Following gene design of individual *recA* protomers, the recoded and modified *recA* genes were prepared synthetically (IDT). Purified genes were restriction enzyme-digested and ligated into pUC19 vectors (New England Biolabs, NEB). Iterative rounds of digestion and ligation were used to produce each successively longer *recA* oligomer construct. The final coding sequences were sub-cloned into pET41 expression vectors (**Fig. S1**). The RecA3x mutants containing either G22Y or G108Y in each protomer were generated using iterative rounds of either site-directed mutagenesis or sub-cloning from synthetic gene fragments. The identity and sequence of each plasmid was confirmed through restriction mapping and sequencing.

### Protein Expression and Purification of RecA constructs

Oligomeric RecA overexpression plasmids were transformed into *recA*-deficient BLR(DE3) cells (Thomas Scientific) to further minimize *in vivo* recombination of expression vectors. Cultures were grown in Luria-Bertani (LB) media (Sigma), induced by addition of 1 mM (final) isopropyl β-D-1-thiogalactopyranoside (IPTG) after culture reached an OD_600_ >1.2, followed by expression for four hours at 37 °C. The cells were then harvested by centrifuging at 3,000 rcf for 20 minutes and resuspended in 10 mL of RecA Lysis Buffer (50 mM sodium phosphate, pH 8.0, 500 mM KCl, 500 mM NaCl, 25 mM imidazole) per gram wet cell pellet with cOmplete EDTA-free protease inhibitor cocktail tablet (Sigma). Resuspended cells were lysed by passing five times through a Microfluidics M-110P microfluidizer. 250 units of benzonase was added per 10 mL lysate and incubated for 30 minutes at 4° C. For each RecA sample, the lysate was clarified by spinning for 15 minutes at 25,000 rcf and the retained supernatant was flowed twice over HisPur Cobalt Resin (Thermo Scientific) pre-equilibrated with RecA Lysis Buffer. The resin was washed with Cobalt Load/Wash Buffer (50 mM sodium phosphate, pH 8.0, 500 mM KCl, 500 mM NaCl, 100 mM imidazole), and the sample was eluted with Cobalt Elution Buffer (50 mM sodium phosphate, pH 8.0, 500 mM NaCl, 400 mM imidazole). Saturated ammonium sulfate solution was added to the eluent at 4 °C to 25% saturation followed by 30 minute equilibration. The solution was centrifuged at 3,000 rcf for 30 minutes and the pellet was discarded. This precipitation step was repeated at 48% ammonium sulfate saturation and the pellet was resuspended in Q Column Load/Wash Buffer (20 mM Tris-HCl, pH 8.0, 20 mM NaCl, 2 mM tris (2-carboxyethyl) phosphine hydrochloride (TCEP)) and then dialyzed against Q Column Load/Wash Buffer for 2 hours at 4° C to remove remaining ammonium sulfate. Using an ÄKTA FPLC system, the sample was run through a HiTrap Q HP 5 mL column (Cytiva). The column was washed with a mixture of 95% Q Column Load/Wash Buffer and 5% Q Column Elution Buffer (20 mM Tris-HCl, pH 8.0, 1.5 M NaCl, 2 mM TCEP) followed by sample elution with a mixture of 85% Q Column Load/Wash Buffer and 15% Q Column Elution Buffer. The eluent was then flowed over a Heparin HiTrap HP 5 mL column (Cytiva) and followed with a wash of a mixture of 85% Q Column Load/Wash Buffer and 15% Q Column Elution Buffer. The sample was eluted from the heparin column with a mixture of 70% Q Column Load/Wash Buffer and 30% Q Column Elution Buffer. Heparin elution peaks were pooled and concentrated using Amicon Ultra Centrifugal Filters, 10K MWCO (Millipore Sigma). The sample was flowed onto a HiLoad 16/600 Superdex 200 pg gel filtration column (Cytiva) equilibrated with RecA Storage Buffer (20 mM Tris-HCl, pH 8.0, 200 mM NaCl, 2 mM TCEP, 10% v/v glycerol) and elution peaks were combined after running on 4-20% SDS-PAGE gels (Bio-Rad) for verification. The protein sample concentration was quantified using Qubit Protein Kit (Thermo) and stored at -80 °C.

### Protein Expression and Purification of LexA, LexA variants, and CI

LexA, LexA variants, and the lambda phage repressor (CI) containing N-terminal polyhistidine tags were cloned in pET41 overexpression plasmids and transformed into BL21(DE3) cells. The acridonylalanine (δ) labeled, C-terminal His-tagged LexA variant (LexA-S119A-Q161δ, termed LexA-δ) was expressed in BL21(DE3) cells containing an auto-inducible plasmid-borne acridonylalanine tRNA synthetase in the presence of 1.0 mM acridonylalanine.^54^ Expressions and purifications of each LexA variant were carried out as previously described,^55^ and an identical protocol was employed for CI. Protein concentrations were quantified using Qubit Protein Kit (Thermo) and protein stocks were stored at -80 °C.

### LexA Labeling with CF488A (LexA-CF)

As LexA has no native Cys residues, a tolerant position was mutated to facilitate making a labeled LexA variant.^56^ The His-LexA W201C mutant was generated using a Q5 Site-Directed Mutagenesis Kit (NEB) and the mutation was confirmed via sequencing. The mutant was expressed and purified in the same way as described for wild type LexA. Purified His-LexA W201C was incubated in LexA Storage Buffer (50 mM Tris-HCl, pH 7.0, 200 mM NaCl, 0.5 mM EDTA, 10% glycerol) and equimolar TCEP for 30 minutes on ice before addition of a 5-fold molar excess of the green fluorescent CF488A maleimide dye (Sigma). The mixture was gently agitated overnight at 4 °C in a 1.5 mL amber tube. The post-labeling reaction was spun down for 15 minutes at 15,000 rcf at 4 °C and supernatant was added to an equal volume of LexA Cobalt Wash Buffer (20 mM sodium phosphate, pH 6.9, 500 mM NaCl, 25 mM imidazole) and loaded onto equilibrated HisPur cobalt resin via batch binding for approximately 2 hours at 4 °C. The flow through was discarded followed by a wash with LexA Cobalt Wash buffer until there was no detectable fluorescence in the wash. The sample was eluted with LexA Cobalt Elution Buffer (20 mM sodium phosphate, pH 6.9, 500 mM NaCl, 400 mM imidazole) twice. The sample was then dialyzed overnight at 4 °C into LexA Storage Buffer and spun down for 15 minutes at 15,000 rcf at 4 °C to remove precipitated protein. The concentration of LexA-CF was determined by both calculating it from the absorbance at 490 nm (ε_490_ = 70,000 M^-1^ cm^-1^ for CF488A dye) and measuring it from an in-gel LexA standard curve. The sample was stored at -80 °C.

### LexA Cleavage Assays

RecA constructs were pre-activated in 1X Activation Buffer (70 mM Tris-HCl, pH 7.5, 150 mM NaCl, 10 mM MgCl_2_, 2 mM TCEP, 0.25 mM ATPγS) for 3 hours at 25 °C with an 18-mer (GGT)_6_ ssDNA substrate containing six GGT repeats. Pre-activated RecA* mixtures were then mixed 1:1 with LexA- or λ CI-containing mixtures and incubated at 25 °C. Zero-minute timepoints were collected by independently adding 5 µL of each mixture to 10 µL 2X Laemmli Buffer (Bio-Rad) and boiling at 95 °C for 5 minutes. Reactions were stopped by adding equal volumes of 2X Laemmli Buffer to the reaction mixtures at the appropriate timepoints. For the qualitative LexA cleavage assays, the final concentrations were as follows: [RecA]_protomer_ = 1 µM, [(GGT)_6_] = 1 µM, [LexA] = 1000 nM. For the qualitative λ phage CI cleavage assays, the final concentrations were as follows: [RecA]_protomer_ = 2 µM, [(GGT)_6_] = 2 µM, [CI] = 2 µM. For the quantitative LexA cleavage assays, the final concentrations were as follows: [RecA]_protomer_ = 1 µM, [(GGT)_6_] = 1 µM, [LexA-CF] = 100 nM. For RecA mutant screening, the final concentrations were as follows: [RecA]_protomer_ = 1 µM, [(GGT)_6_] = 1 µM, [LexA]_unlabeled_ = 1900 nM, [LexA-CF] = 100 nM. As a negative control, RecA Storage Buffer was used in an activation reaction in lieu of RecA protein. The standard curve using Coomassie stain and CF488A dye were used to establish the linear range of quantification for the assay (**Fig. S2**). Following reaction stoppage, 10 µL of each timepoint was loaded onto a 15% SDS-PAGE gel. Reactions containing LexA-CF variant were immediately imaged on a Typhoon imager using the Cy2 laser line at a PMT voltage of 250 V. Qualitative gels were Coomassie stained and imaged on a Bio-Rad Gel Doc XR+. For ATP/ssDNA-dependence qualitative assays, either ssDNA, ATPγS, or both were withheld. For noted samples, RecA working stocks were pre-mixed with 250 units of benzonase (Sigma) and incubated for 20 minutes at 25 °C. Gel band intensities were quantified using ImageJ. Percent cleavage was calculated by ratiometric comparison of background-corrected cleavage band intensity to background-corrected full-length band intensity at each time point according to the following equation:

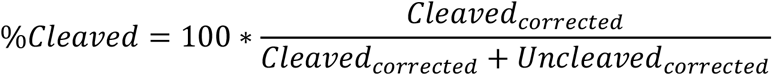

### RecA Filamentation Assay

RecA constructs at various concentrations were activated as described in the LexA cleavage assay, with 2 nM 3’-FAM labeled (GGT)_6_ oligonucleotide (IDT) and 70 µg/mL BSA. The samples were transferred to Kimble 6×50 mm borosilicate glass culture tubes after a 3 hour incubation and spun down in a tabletop microfuge. Anisotropy measurements were taken on a Panvera Beacon2000 with a 490 nm excitation filter and a 525 nm emission filter (Farrand Controls). Blank solutions were prepared with no labeled (GGT)_6_, and control solutions were prepared with no RecA protein. Binding affinities, baseline anisotropy, and change in anisotropy were determined using GraphPad Prism 9.3.1 non-linear regression fitting to the quadratic equation with a fixed (GGT)_6_ (Oligo) concentration of 2 nM:

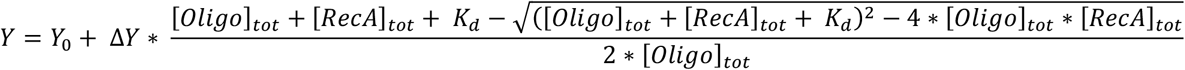

Here Y is the anisotropy at a given concentration of RecA, Y_0_ is the baseline anisotropy, and ΔY is the change in anisotropy from baseline to the highest value. Since RecA WT is known to be a cooperative binder to ssDNA, an Adair model was used for fitting WT data, with two K_assoc_ values defined for nucleation (K_1_) and filament extension (K_2_) for up to six RecA unit binding events:

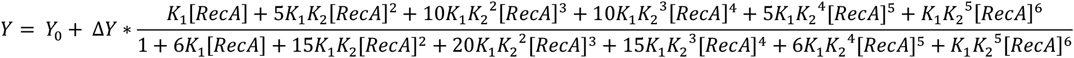

### LexA Binding Assay

RecA constructs were serially diluted and then activated as described in the LexA cleavage assay, with 70 µg/mL BSA present in the activation buffer. For RecA WT each condition had equimolar (GGT)_6_ oligonucleotide, while for RecA3x and RecA4x there was 40 µM (GGT)_6_. These concentrations maximized the amount of filamented RecA while avoiding inhibition of RecA WT filament extension via large excess of ssDNA. Pre-activated filaments were incubated for 2 hours at 25 °C, after which LexA-δ was added and the solution was further incubated for 30 min. The anisotropy for each sample was measured on the Panvera Beacon2000 with a 387 ± 11 nm excitation filter and 448 ± 20 nm emission filter (Edmond Optics). Binding affinities, baseline anisotropy, and change in anisotropy were again determined using GraphPad Prism 9.3.1 non-linear regression fitting to the previous quadratic equation with [LexA-δ] in place of [(GGT)_6_], fixed at 50 nM.

### Steady-State Kinetics of LexA Cleavage

For steady-state kinetic experiments, the concentration of RecA* was reduced in order to satisfy the steady-state requirement of excess substrate (LexA) over ‘enzyme’ (RecA*), and the DNA binding affinities were used to estimate the amount of active RecA* in solution under each reaction condition. RecA protomer concentrations in this assay were 600 nM for RecA3x and 450 nM for both RecA WT and RecA4x, with equimolar (GGT)_6_. Pre-activation and cleavage reactions were carried out as previously described, except that each reaction contained a fixed amount of LexA-CF (100 nM) and increasing amounts of unlabeled LexA for a total [LexA] ranging from 310 nM to 10 µM. Rates of cleavage were determined for early timepoints to ensure the linear phase of the reaction was captured, and initial rates were plotted against [LexA]_total_ in GraphPad Prism 9.3.1. Data were fit to the standard Michaelis-Menten equation using nonlinear regression. The V_max_ value was then normalized using the estimated [RecA*] to yield a k_cat_, from which we also calculated a k_cat_ / K_M_.

## RESULTS AND DISCUSSION

### Design of oligomeric RecA constructs

Prior work aiming to define the SOS activation complex exclusively relied upon studying monomeric RecA (hereafter called RecA WT). We considered the possibility that oligomeric RecA constructs, capable of forming defined filament sizes, could be employed to offer new insights into LexA engagement. To this end, we based our design on the linked oligomer strategy used to solve the structure of the RecA* filament. In this approach, the Pavletich group used translationally fused, end-capped constructs between four to six RecA units long. This strategy was also used to make a two unit construct in order to probe for the minimal requirements necessary for homologous recombination.^52,53^ Our general design revolved around linking a defined number of RecA protomers via a 14-amino acid linker in a single open reading frame (**Fig. 1B**).^52^ In order to prevent further oligomerization on either end, the N-terminal RecA unit contained a deletion of amino acids 1-30, which removes the α-β oligomerization motif, and the C-terminal RecA unit possessed the oligomerization disrupting mutations C117M, S118V, and Q119R (**Fig. 1B**).^57^ In addition, each internal protomer contained a truncation of the C-terminal-most amino acids 336-353, which removes an allosteric regulatory region and has been shown to increase DNA-binding capabilities of RecA.^58^ We extended this approach to generate RecA oligomers containing from one (RecA1x) to six (RecA6x) RecA protomers. In our design, we further performed recoding of the individual RecA protomers to facilitate downstream mutagenesis and to mitigate *in vivo* recombination that was observed with higher order constructs. In the application of our design through construct expression and purification, we achieved a high degree of purity for oligomers one to four units long and a moderate degree of purity for oligomers five to six units long (**Fig. 1C**).

### Engineered RecA constructs are active for LexA cleavage

Although a subset of these RecA species were previously studied in the context of homologous recombination, we were interested in their capacity for SOS activation. As SOS activation is ultimately achieved through cleavage of LexA, we reasoned that LexA cleavage activity could serve as a proxy for the preceding biochemical events. By focusing on LexA cleavage first, we could then determine which of the RecA constructs to use for downstream analysis of the individual biochemical steps in SOS activation. For an initial qualitative assessment of cleavage, we utilized conditions in which monomeric RecA WT can be activated to form RecA* and promote LexA cleavage (**Fig. 2A**). For increased sensitivity and more rigorous quantification, we also generated a LexA W201C mutant and labeled it with a fluorescent CF488A maleimide dye (LexA-CF). This variant allows us to accurately quantify small changes in LexA cleavage when spiked into the sample without significantly affecting the overall auto-proteolysis rates (**Fig. S3-S4, Fig. 2B**). RecA WT was included as a control to establish a baseline of LexA cleavage under these conditions. Consistent with the fact that RecA1x is incapable of forming stable oligomers, we observed no acceleration of LexA cleavage over the baseline (**Fig. 2B**). We did, however, observe detectable cleavage over the baseline with RecA2x, reaching 8 ± 1 % cleavage after 60 minutes (**Fig. 2B**). Cleavage increased substantially with RecA3x (32 ± 2 %), and then increased in a stepwise fashion with each progressively longer RecA construct. Notably, RecA6x displayed the highest level of LexA cleavage of the engineered constructs (82 ± 1 %), which represented the maximal amount of cleavage detectable in this assay and was equivalent to the activity of RecA WT (**Fig. 2B**).

**Figure 2.**
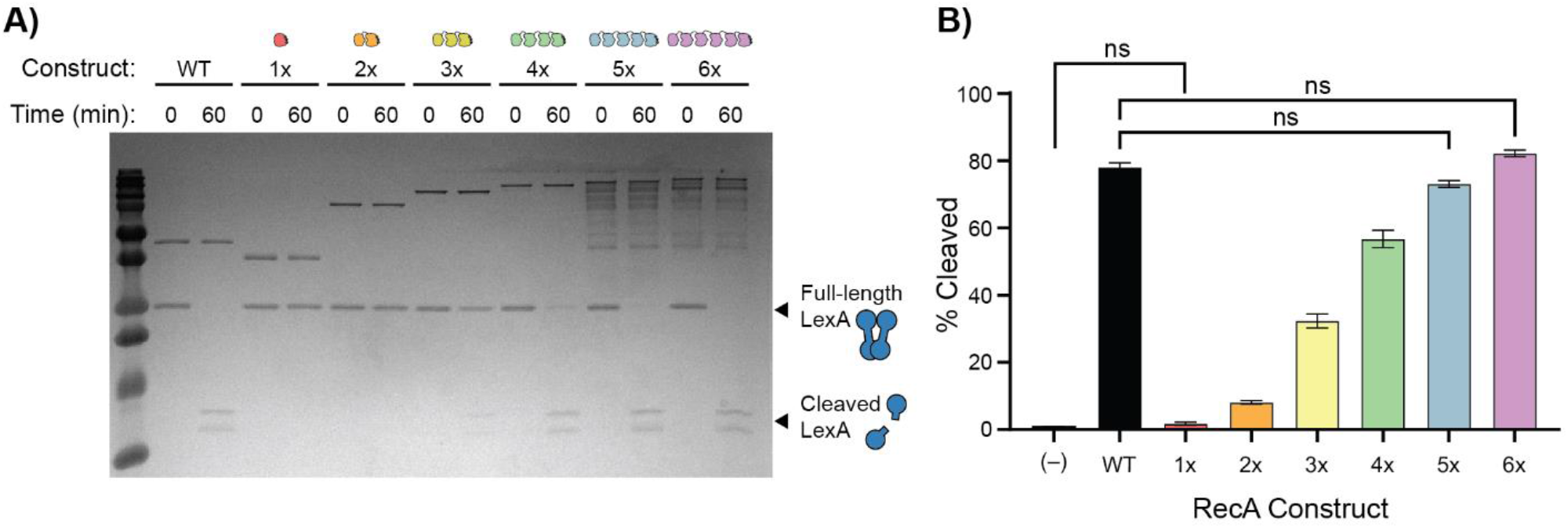
RecA*-stimulated LexA cleavage. **A)** RecA*-stimulated LexA cleavage assay run for 60 minutes and visualized on a 15% SDS-PAGE gel after Coomassie staining. Each reaction contained 1 µM each of RecA_protomer_, (GGT)_6_ ssDNA oligonucleotide, and LexA. **B)** Quantification of fluorescent LexA-CF cleavage reaction by RecA* constructs. These reactions were performed in triplicate under similar conditions as in **Fig. 2A** with LexA-CF allowing for rigorous quantification. Data shows the means and standard deviation of percent cleavage. The cleavage efficiency of each construct was compared pairwise using one-way ANOVA assuming non-uniform standard deviations. All comparisons were significant at *p* < 0.05 unless otherwise noted (ns).

The observation that RecA2x increased cleavage above baseline was intriguing. Prior work has suggested that RecA2x construct does not form a stable extended filament, but can support transient and weak DNA associations that can support homologous recombination.^53^ Given the increase in cleavage observed with RecA3x, our data suggests a role for RecA units in addition to the 2x interface. To test the generalizability of the posited “minimal” RecA requirement, we also examined the ability of RecA2x and RecA3x to catalyze the auto-proteolysis of another RecA co-proteolytic substrate – the λ phage repressor (CI), which has a similar catalytic core and auto-proteolytic mechanism to that of LexA.^59-61^ With CI, we observed no cleavage above baseline with the RecA2x construct, and a substantial increase in cleavage proficiency using RecA3x (**Fig. S5**). The consistent pattern observed suggests that a RecA3x interface can efficiently engage LexA and that this interface may be shared by RecA’s other substrates that share a similar mode of activation (**Fig. S5**).

In prior work, experiments that utilized chemical crosslinking yielded a model of seven RecA protomers per LexA whereas cryo-EM studies proposed a more simplistic 2-3 RecA units per LexA interaction ratio.^48,50^ Our data better align with the prior low resolution structural work, given the proficiency of RecA3x for LexA cleavage. Beyond these preliminary insights, to further probe the mechanism underlying RecA*-LexA complex formation, we next aimed to explore the proficiency of each individual biochemical step leading up to LexA cleavage, starting with formation of RecA* filaments on ssDNA.

### Longer engineered RecA constructs are competent for filament formation

When forming nucleoprotein filaments on ssDNA, RecA WT is known to be cooperative, leading to a slow nucleation phase followed by rapid filament extension.^43,46,62,63^ As noted earlier, it has been observed that although RecA2x may be capable of binding ssDNA, it does so at a reduced capacity relative to RecA WT.^53^ Meanwhile, RecA4x and larger have been shown to be capable of binding to ssDNA stably enough for structural studies via X-ray crystallography.^52^ We therefore anticipated a range of ssDNA-binding activity with our RecA series and posited that filament formation could be contributing to the observed pattern of LexA cleavage.

In order to track filament formation, we utilized a 3’-FAM labeled ssDNA oligonucleotide in a fluorescence anisotropy-based assay. By measuring the increase in fluorescence anisotropy as a function of added RecA at equilibrium, we can model the interaction to determine apparent binding affinities for each of the constructs (**Fig. 3A**). Notably, formation of RecA* is dynamic, requiring first nucleation on ssDNA followed by filament extension, a process during which filaments may be subject to destabilization or filament rearrangement.^43,64-66^ To ensure that our measurements are indeed taken at equilibrium and that filaments are not subject to stochastic destabilization under our conditions, we performed a filamentation time course with either RecA WT or RecA4x. This experiment showed that equilibrium is achieved by 1 hour, prompting our use of a consistent 3-hour incubation for our equilibrium measurements (**Fig. S6**).

**Figure 3.**
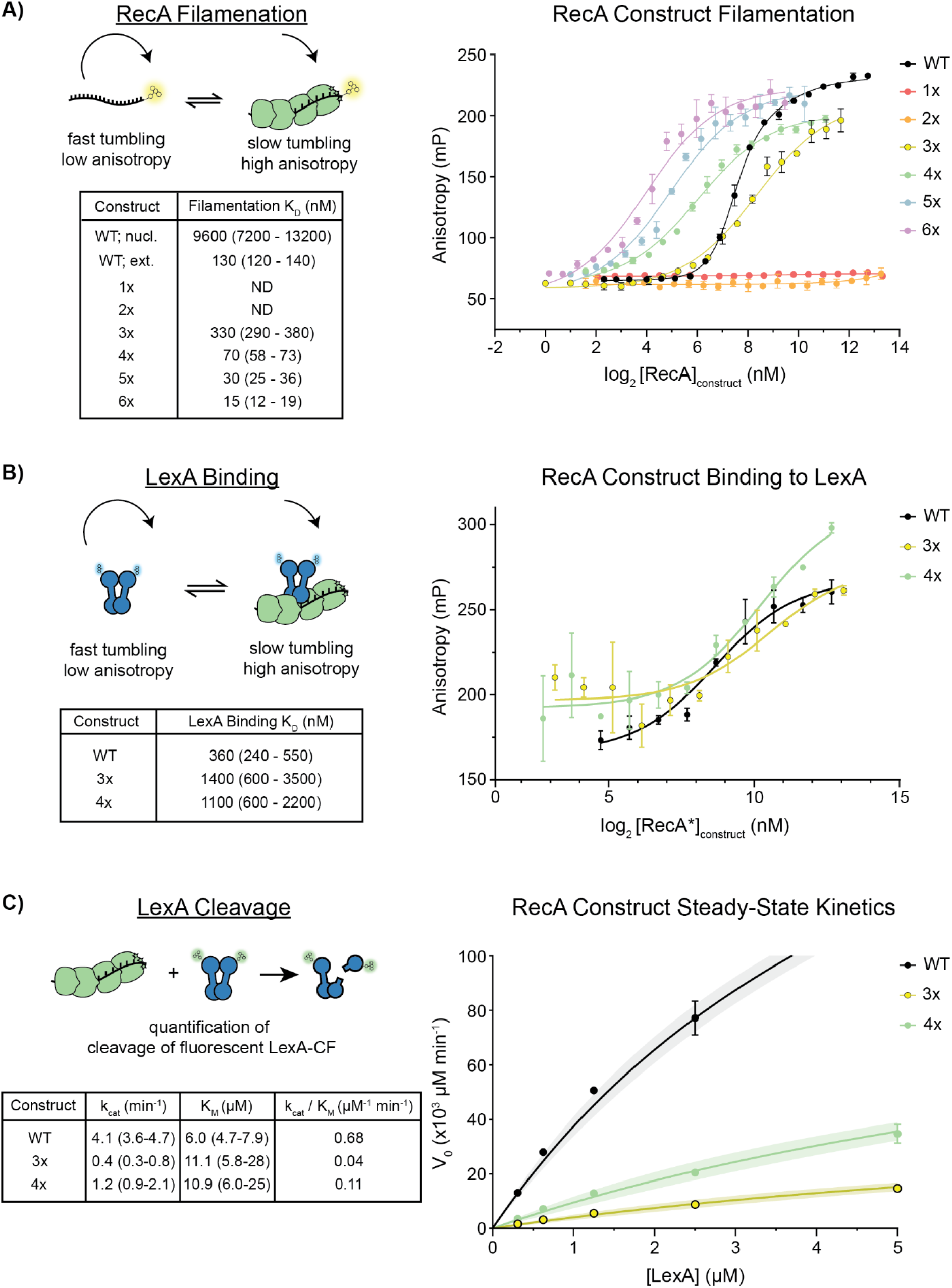
Quantitative biochemical assessment of SOS activation steps for RecA constructs. **A) RecA Filamentation.** Filament formation is tracked by anisotropy using fluorescent end-labeled ssDNA. Means and standard deviations of anisotropy measurements are plotted as a function of [RecA]_construct_ from three replicates (on the right), with data fit to either the quadratic binding model or Adair model of cooperative binding (solid lines). The associated best-fit K_D_ value of each construct and the 95% confidence interval are given as the filamentation K_D_ (table). **B) LexA Binding**. Complex formation is tracked by anisotropy with fluorescent, cleavage-incompetent LexA-δ. Means and standard deviations of anisotropy measurements are plotted as a function of [RecA*] constructs from three replicates (on the right), with data fit to a quadratic binding model (solid lines). Best-fit binding affinities and 95% confidence intervals are given (table). **C) LexA Cleavage**. LexA cleavage is tracked by quantification of cleavage product using fluorescent labeled LexA-CF. Means and standard deviations for initial velocities of the reaction are plotted as a function of [LexA] for each RecA construct from three replicates (on the right), with the data fit to the Michaelis-Menten equation (solid lines) with 95% confidence intervals shown (shaded regions). Derived values for k_cat_ and K_M_, the associated 95% confidence intervals, and calculated specificity constants are shown (table).

As expected, the RecA1x construct shows no appreciable ssDNA binding even at high concentrations. RecA2x also appears to have severely impaired ssDNA binding properties comparable to RecA1x (**Fig. 3A**). As previously discussed, it has been demonstrated that the RecA2x construct is catalytically proficient at homologous recombination despite not forming traditional filaments. Together these results lend further support to the idea that RecA2x is forming transient nuclei on ssDNA rather than stable filaments.^53^

RecA WT and each of the constructs from RecA3x to RecA6x showed proficient RecA* filament formation as detected by a change in anisotropy at equilibrium. In comparing the results of RecA WT to the linked oligomeric constructs, the concentration-dependent filament formation curves appeared different. With monomeric RecA WT, the binding curve indicated a high degree of cooperativity, in agreement with previous findings (**Fig. 3A**).^27,62^ Interestingly, cooperativity was not evident for the engineered construct binding curves. This result likely reflects one potential advantage of studying engineered constructs in the complex pathway of SOS activation, as filamentation can be reduced to an apparent bimolecular association as opposed to cooperative biphasic filament extension.

Because of the differences in observed cooperativity, we initially fit RecA WT to a cooperative binding model while we treated the oligomeric linked constructs as bimolecular associations. Using this approach, a clear pattern emerged of increasing affinity for ssDNA with an increasing number of protomers (**Fig. 3A**). The effect was non-linear, with the affinity of RecA3x nearly 25-fold weaker than that of RecA6x. We speculate that differences in affinity reflect slower dissociation rates with the higher number of protomers engaged with ssDNA. While protomer cooperativity does not manifest in the shape of overall binding curves, such cooperative effects could be driving the higher apparent binding affinity the larger constructs have for ssDNA.

Although we observed clear differences in ssDNA binding affinity, it was not clear if these differences would manifest in different levels of [RecA*] at the conditions used to study LexA cleavage. When using the derived binding affinity along with our known concentrations in the LexA cleavage assay ([RecA]_protomer_ = 1 µM, [(GGT)_6_] = 1 µM), we found that the anticipated amounts of activated [RecA*] filament were similar (range from 160 – 230 nM) and not able to account for the differences in cleavage that we had observed (**Fig. 3A, Fig. S7**). We further recognized that the relevant species for modeling binding could either involve the RecA construct or total RecA protomers. For example, 200 nM of RecA3x would be equivalent to 600 nM of RecA protomer. However, whether these curves were modeled using total protomer concentrations or construct concentrations, the amount of interpolated [RecA*] under our qualitative cleavage reaction conditions was comparable (**Fig. S7**).

Because the differences in filamentation competencies between engineered constructs did not sufficiently account for the range in LexA cleavage activity we observed, we next focused on the subsequent biochemical step in SOS activation, LexA binding. We chose to focus downstream analysis on RecA3x and RecA4x, as these two constructs (1) are active for LexA cleavage albeit to different degrees, (2) demonstrate the ability to bind ssDNA, and (3) can be reliably expressed and purified in large quantities.

### Oligomeric RecA* binds more weakly to LexA

Given that both RecA3x and RecA4x promoted LexA cleavage at rates slower than RecA WT, we posited that either LexA binding or allosteric co-protease kinetics might be impacted in the modified constructs. Our lab has previously employed fluorescently-labeled LexA variants to show that much of the DNA binding NTD of LexA is dispensable for RecA* binding.^55^ We built on this precedent to study the binding of LexA to RecA oligomers. In order to isolate the binding step and prevent downstream proteolysis, we used LexA with an S119A mutation that removes the catalytic serine responsible for autoproteolysis. We also used the unnatural amino acid acridonylalanine (Acd, δ) and amber suppression to incorporate Acd as a minimally-perturbing fluorescent probe into LexA at position Q161 (LexA S119A Q161δ, LexA-δ).^55,56^ The long fluorescence lifetime of Acd (15 ns) offers a large dynamic signal range, thus allowing us to study complex formation by fluorescence anisotropy (**Fig. 3B**). Unlike with RecA WT, where a large excess of ssDNA has been observed to be inhibitory for filament formation, the fusion of RecA protomers together in our constructs allowed us to drive filament formation allowing us to measure anisotropy changes as a function of [RecA*].^67-70^

Using this approach, with RecA WT, LexA-δ anisotropy increases with [RecA*] revealing a bimolecular binding pattern with a measured affinity of 360 nM. Activated RecA3x and RecA4x showed similar patterns to one another overall, with K_D_ values of 1.4 µM and 1.1 µM for RecA4x and RecA3x, respectively. This result suggests that these constructs are competent for LexA binding, albeit with slightly weaker affinities than that of activated filaments generated by RecA WT (**Fig. 3B**). The tighter binding affinity of RecA WT may be a function of the fact that native RecA can form more “perfect” filaments on the ssDNA oligomer due to its ability to rearrange and bind unit-by-unit, whereas the oligomeric RecA constructs could potentially bind form “imperfect” filaments. Alternatively, it is possible that the differences in the number of “docking” sites for LexA, which could bind in various registers in the filaments formed by RecA WT, could account for tighter binding. A similar rationale could explain the small decrease in K_D_ of the activated RecA4x relative to RecA3x.

### Steady-state kinetics inform a model of RecA*-LexA engagement

Our results so far provided valuable thermodynamic insight into the stability of RecA* formation by each oligomeric RecA species and each construct’s capacity for LexA binding. We next aimed to define the steady-state kinetic parameters to discriminate between enzyme turnover, k_cat_, and the balance between complex formation and product release, K_M,_ in LexA cleavage. Defining these parameters can support future efforts to yield stable complexes for structural characterization or help with exploring the effects of targeted mutations on individual steps in SOS activation.

To this end, we mixed a low nanomolar amount of either activated RecA WT, RecA3x, or RecA4x with an excess of LexA and monitored the cleavage over the linear range of product formation (0-15 minutes). In this experimental setup, although LexA is capable of auto-proteolysis, we considered the activated RecA species as an “enzymatic” species, given its critical co-catalytic role. To increase sensitivity and reduce noise in quantitative analysis, we again employed the fluorescent LexA-CF as a spike-in with LexA WT and collected serial time points in cleavage reactions (**Fig. 3C, Fig. S8**). Fitting the initial rates of cleavage against concentrations of total LexA to Michaelis-Menten models yielded the steady-state kinetic values, k_cat_ and K_M_, for each RecA species. We observed that RecA WT has about a 3-fold higher k_cat_ and 2-fold lower K_M_ as compared to RecA4x, which results in an overall 6-fold improvement of the specificity constant (k_cat_ / K_M_) for RecA WT over RecA4x (**Fig. 3C, Fig. S9**). While we observe a similar 2-fold change in K_M_, the impact on k_cat_ is more significant with RecA3x compared to RecA WT, resulting in an overall 20-fold difference in the specificity constant between RecA WT over RecA3x (**Fig. 3C, Fig. S9**).

Integrating across the biochemical observations we describe above, which parse each separate step in SOS activation with distinct fluorescence-based assays, provides a more comprehensive model for the essential constituents involved in the formation of the SOS activation complex. We demonstrate that inefficient formation of stable RecA* filaments with RecA1x and RecA2x is associated with an impaired ability to support efficient LexA autoproteolysis. Filament formation appears bimolecular for the longer linked RecA constructs (RecA3x through RecA6x), as opposed to the cooperative binding observed for monomeric RecA WT. For the longer linked RecA constructs, a positive non-linear relationship exists between binding parameters and the number of oligomeric units, suggesting that synergistic binding of monomers drives stable RecA* formation. Nonetheless, RecA* formation alone does not explain differences in the overall rate of LexA cleavage. Both RecA3x* and RecA4x* show modestly lower binding affinities for LexA, and these differences extend to higher K_M_ values for cleavable LexA as well. The overall decrease in catalytic efficiency (k_cat_/K_M_) for both RecA3x and RecA4x is primarily a result of depressed k_cat_, which suggests that inherent properties of the engineered constructs, such as the unnatural linkers or the absence of accessory RecA subdomains, might impact co-protease function. When we compare the RecA3x and RecA4x constructs, both are readily able to support LexA cleavage; however, the higher catalytic efficiency of RecA4x could be attributed to the additional RecA unit either offering improved chances of binding in a mode supportive of productive catalysis or directly making additional contacts with LexA.

Having demonstrated the proficiency of oligomeric RecA constructs in supporting LexA activation, we next aimed to leverage our experimental system to address targeted questions regarding the nature of the SOS activation complex, including the viability of existing models for engagement of LexA and RecA*.

### ATP and DNA are both strictly required for LexA co-proteolytic activity

It has been shown that the ATP and DNA binding activities of RecA are indispensable for both cell viability under genotoxic stress and *in vivo* biochemical competence of homologous recombination.^49^ However, it has not been directly demonstrated whether filamentation, and the resulting conformational extension of RecA, is strictly necessary for the co-protease activity of RecA or if colocalization of several RecA protomers alone is capable of engaging LexA. For example, alterations at Pro67 of RecA that relax nucleotide cofactor specificity can lead to either highly constitutive co-protease activity or activity requiring significantly shorter ssDNA cofactors, suggesting that RecA oligomers may not need to adopt the specific conformation of the extended, active filament bound to ATP.^71^ Additionally, LexA cleavage experiments performed with high concentrations of crowding reagents, such as PEG, have been shown to stimulate LexA cleavage even in the absence of ssDNA.^72,73^

Because each RecA protomer in our constructs is translationally fused, we next aimed to test whether ATP and DNA are obligate parts of the minimally essential activated form of RecA for LexA activation, or whether colocalization of RecA subunits in the linked proteins is sufficient to interface with LexA (**Fig. 4A**). To explore the requirements for active RecA, we systematically tested the RecA3x construct to determine whether co-proteolytic activity was retained in conditions that withheld either ATPγS, ssDNA, or both. To control for the possibility that trace nucleic acid was carried over from purification, one of the samples lacking ssDNA was pre-treated with benzonase before addition to LexA. As expected, RecA WT did not show activity when either ATPγS or ssDNA was withheld. Interestingly, RecA3x followed the same pattern as RecA WT. The linked RecA3x was unable to support LexA cleavage by itself, and activity was not elicited by either binding to ATPγS alone or ssDNA alone. As demonstrated in the earlier analysis of the various RecA oligomeric constructs, in the presence of both ATPγS and ssDNA, RecA3x was proficient in LexA cleavage (**Fig. 4B**).

**Figure 4.**
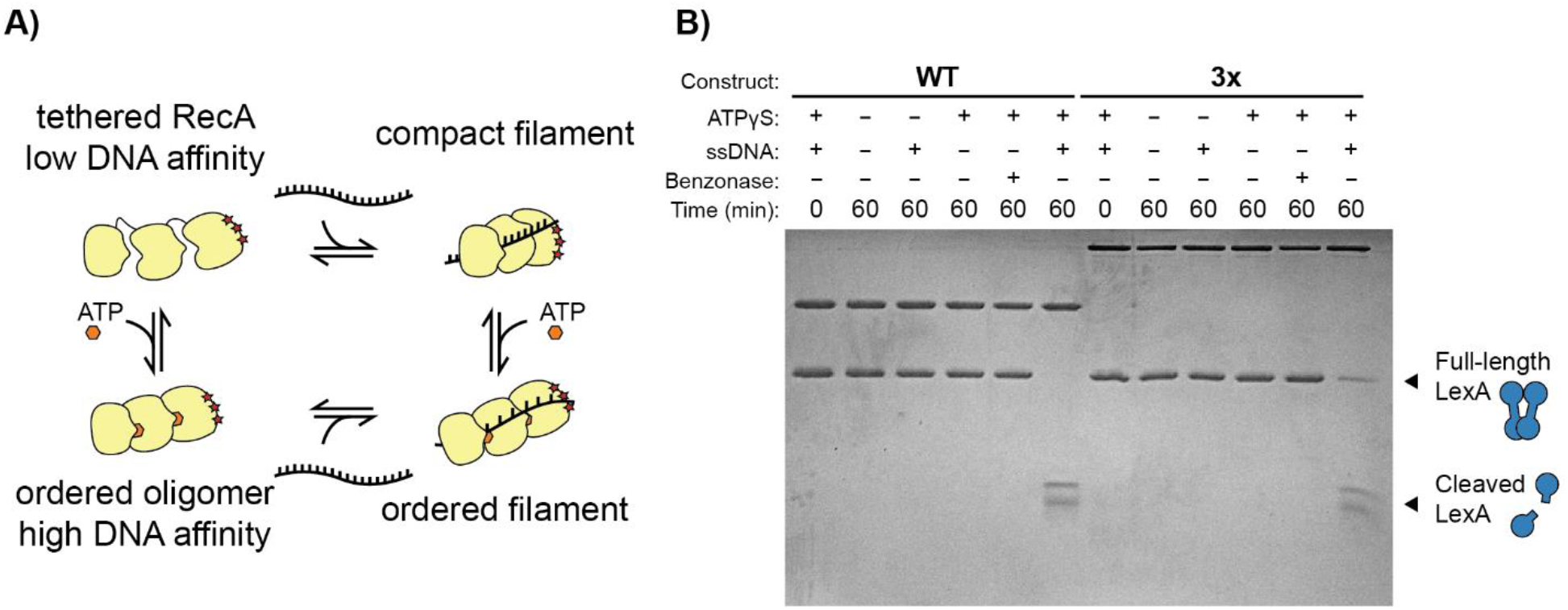
ATP and ssDNA-dependence of RecA constructs. **A)** Graphical representation showing the potential forms of the engineered RecA constructs in the presence or absence of either ATP or DNA. **B)** Qualitative LexA cleavage assays as in **Fig. 2A**, with either ATP, DNA, or both components withheld, visualized on a 15% SDS-PAGE gel after Coomassie staining. Benzonase was added as an additional control against trace nucleic acid carryover from purification.

The finding that RecA3x requires both the ATP analog and ssDNA provides biochemical evidence in favor of the extended, helical conformation of RecA* as the biologically active species for SOS activation. This result is consistent with earlier mutagenesis studies that sought to define the dispensability of various biochemical activities of RecA for its biological functions. For example, the *recA*142 mutant is defective in its ability to compete with single stranded binding protein (SSB) for ssDNA, yet it retains its ability to self-polymerize in the absence of DNA.^28^ Due to the limited availability of ssDNA in the cell, the inability of *recA*142 to form RecA* *in vivo* was thought to explain the phenotypic defect in both homologous recombination and SOS activation.^45,74^ Similarly, the dependence of *in vitro* RecA catalytic activities on first forming extended nucleofilaments indicates that although the physiological activities of RecA may diverge, they are subject to the same activation conditions and likely also the same regulatory control.

### Analysis of RecA mutants *in vitro* challenge a proposed model for LexA engagement

Previous efforts to model SOS activation have sought to identify residues on either RecA or LexA that are important for RecA-dependent LexA cleavage. Initial unbiased attempts to identify mutations that are specific to LexA co-proteolysis were unable to find mutations that only impact LexA cleavage without also impacting the other enzymatic activities of RecA.^45^ In more recent work, to bypass the issues of unbiased mutational screening, an evolutionary trace approach was applied to identify different active site “patches” on the basis of their evolutionary importance.^49^ Guided by these identified patches of residues, the authors evaluated individual and combined mutations with *in vivo* assays for LexA cleavage after UV exposure as well as assays for homologous recombination competence. The authors proposed a model where the RecA-LexA interaction interface is formed by seven RecA protomers in the RecA* filament whereby LexA interacts with G22 on RecA_n_ and G108 on RecA_n+6_.^49^ The authors report that mutation of either of these residues to Tyr leads to a loss of LexA cleavage activity *in vivo*, reportedly with only a partial reduction in homologous recombination efficiency compared to WT.

Our findings that RecA3x is active in LexA binding and cleavage do not support a model in which seven RecA protomers are required to form the LexA binding interface. One important advantage of our engineered RecA constructs is that individual mutations can be introduced on a protomer-by-protomer basis, whereas any mutation made to RecA WT will be present throughout the filament. In this way, our system is positioned to address more nuanced questions about specific residues at precise locations within a RecA filament of defined size.

Although our ultimate goal was to tease apart the roles that these residues may play at specific locations along the filament, we were surprised to discover that when tested *in vitro*, the G22Y mutation was indeed active for LexA cleavage in both RecA WT or RecA3x (**Fig. S10**). Similarly, while G108Y led to some decrease in activity, this variant also was able to support LexA cleavage with both the RecA WT and RecA3x constructs (**Fig. S10**). Together, the demonstration that RecA3x is active for LexA cleavage and our *in vitro* data contradicting the proposed importance of certain contact residues provides added evidence for the inadequacy of the proposed model of the SOS activation complex from evolutionary trace approaches. Furthermore, our results highlight how the minimal SOS system and the assays we have developed stand as valuable tools for validating and potentially identifying future mutation candidates in the continued search for the binding interface between RecA* and LexA.

### Summary of minimal model and potential applications

In summary, systematic evaluation has shown the degree to which each of our engineered RecA constructs is capable of the individual steps of SOS activation: filamentation, LexA binding, and LexA cleavage. Notably, the biochemical data from our RecA3x construct support a model in which three consecutive RecA protomers form a minimal active site capable of undergoing filamentation, binding LexA, and catalyzing LexA cleavage. Our proposal that smaller RecA filaments can promote LexA cleavage is in opposition to several previous models of LexA binding, which instead have proposed an interaction interface involving seven consecutive RecA protomers. Furthermore, the design and validation of our system significantly reduces biochemical complexity. As a result, we were able to ask more precise questions about the nature of SOS activation. First, we directly probed the biochemical requirement for ATP and DNA, showing that both components involved in the fully extended RecA filament are necessary for LexA cleavage. Second, we were able to demonstrate that residues previously thought to be critical for LexA binding and cleavage (i.e., G22 and G108) were dispensable for LexA cleavage when evaluating activity *in vitro*.

The study of macromolecular complexes remains challenging in structural biology and biochemistry due to their complex order of assembly and many constituent parts. Systematic evaluation rapidly becomes challenging, even in cases where much is already understood about parts of the complex. Here, we designed and leveraged engineered protein constructs that could reduce this level of complexity. While this work highlights the limitations of previous models for the SOS activation complex, it also establishes a biochemical foundation for future studies. Specifically, with an improved understanding of the essential components necessary for RecA*-LexA complex formation, RecA3x or RecA4x could prove useful in the structural modeling by biophysical approaches such as cryo-electron microscopy or crystallography. Furthermore, the ability to introduce targeted mutations into individual protomers without those mutations being present in every unit represents a distinct advantage of the linked oligomeric species. The possibility of introducing probes that are both protomer-specific and residue-specific raises the possibility of using fluorescence methods such as FRET to characterize the protein-protein interface between LexA and RecA (**Fig. 5**). Results from these studies could greatly inform ongoing efforts to target the SOS activation complex at a molecular level, given the demonstrated potential for small molecule antagonists to potentiate our antibiotic arsenal and slow the acquisition of antibiotic resistance.^75-77^

**Figure 5.**
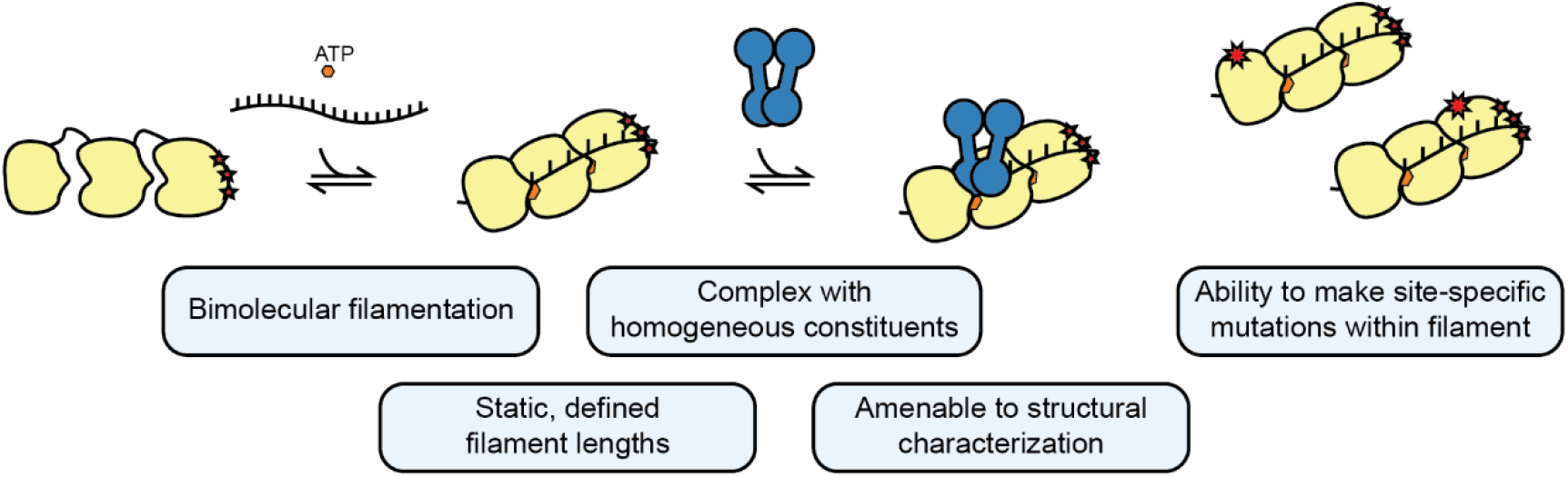
Advantages of engineered RecA constructs. Scheme depicting the kinetic steps of SOS activation. Advantages of the engineered RecA constructs as molecular tools are highlighted.

