## Supplemental Information for "Engineered RecA constructs reveal the minimal SOS activation complex"

**Supplementary Figure 1.** Cartoon schematic of RecA6x plasmid design................................2

**Supplementary Figure 2.** Standard curves for quantitative cleavage assays............................3

**Supplementary Figure 3.** Label on LexA-CF does not affect autoproteolysis.........................4

**Supplementary Figure 4.** Quantitative cleavage of LexA-CF.................................................5

**Supplementary Figure 5.** RecA-stimulated auto-proteolysis of either LexA WT or λ phage repressor (CI). .........................................................................................................................5

**Supplementary Figure 6.** RecA filamentation time course .....................................................6

**Supplementary Figure 7.** Extended analysis of RecA filamentation .......................................7-8

**Supplementary Figure 8.** Representative steady-state LexA cleavage gels.............................8

**Supplementary Figure 9.** Full steady-state kinetic curves for RecA constructs .......................9

**Supplementary Figure 10.** LexA cleavage assessment for RecA mutants...............................10

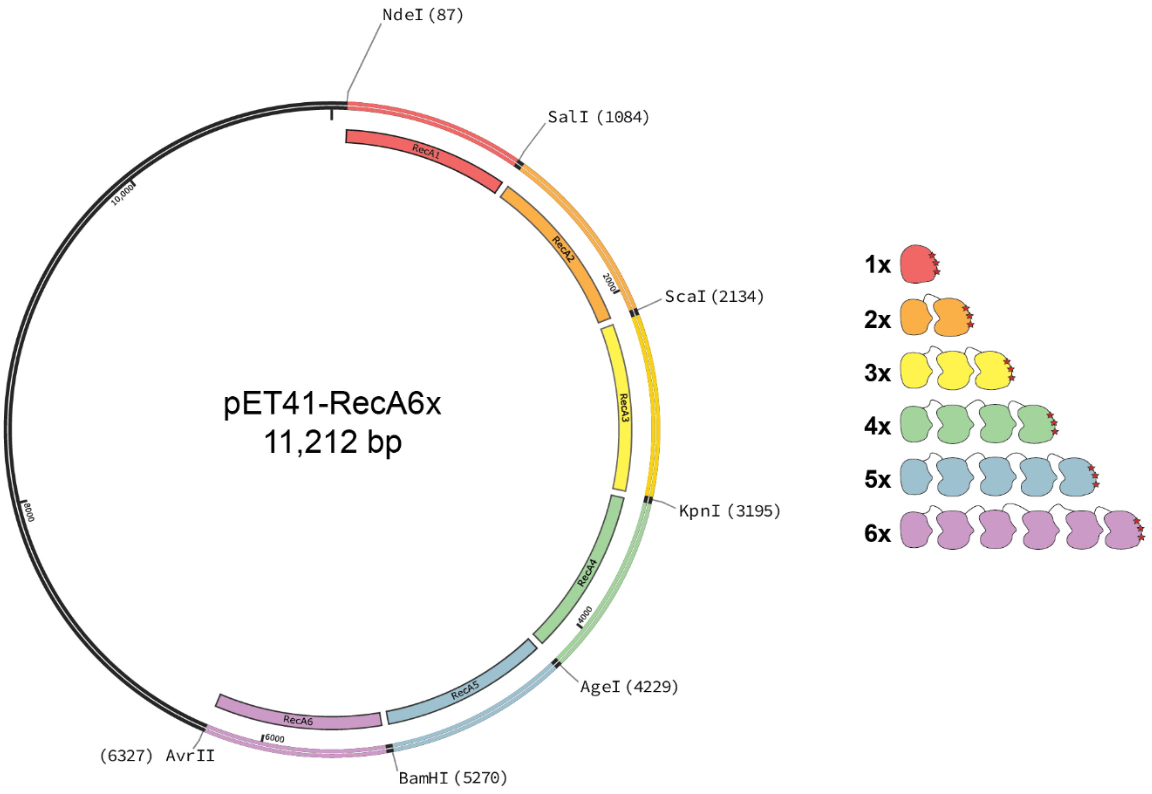


**Figure S1. Cartoon schematic of RecA6x plasmid design.** Scheme of the RecA6x expression plasmid highlights intra-construct restriction sites between each protomer gene used for cloning purposes. Each individual protomer was also genetically recoded to minimize recombination and facilitate downstream mutagenesis. At right is the color coding of the cartoon schematic of RecA constructs.

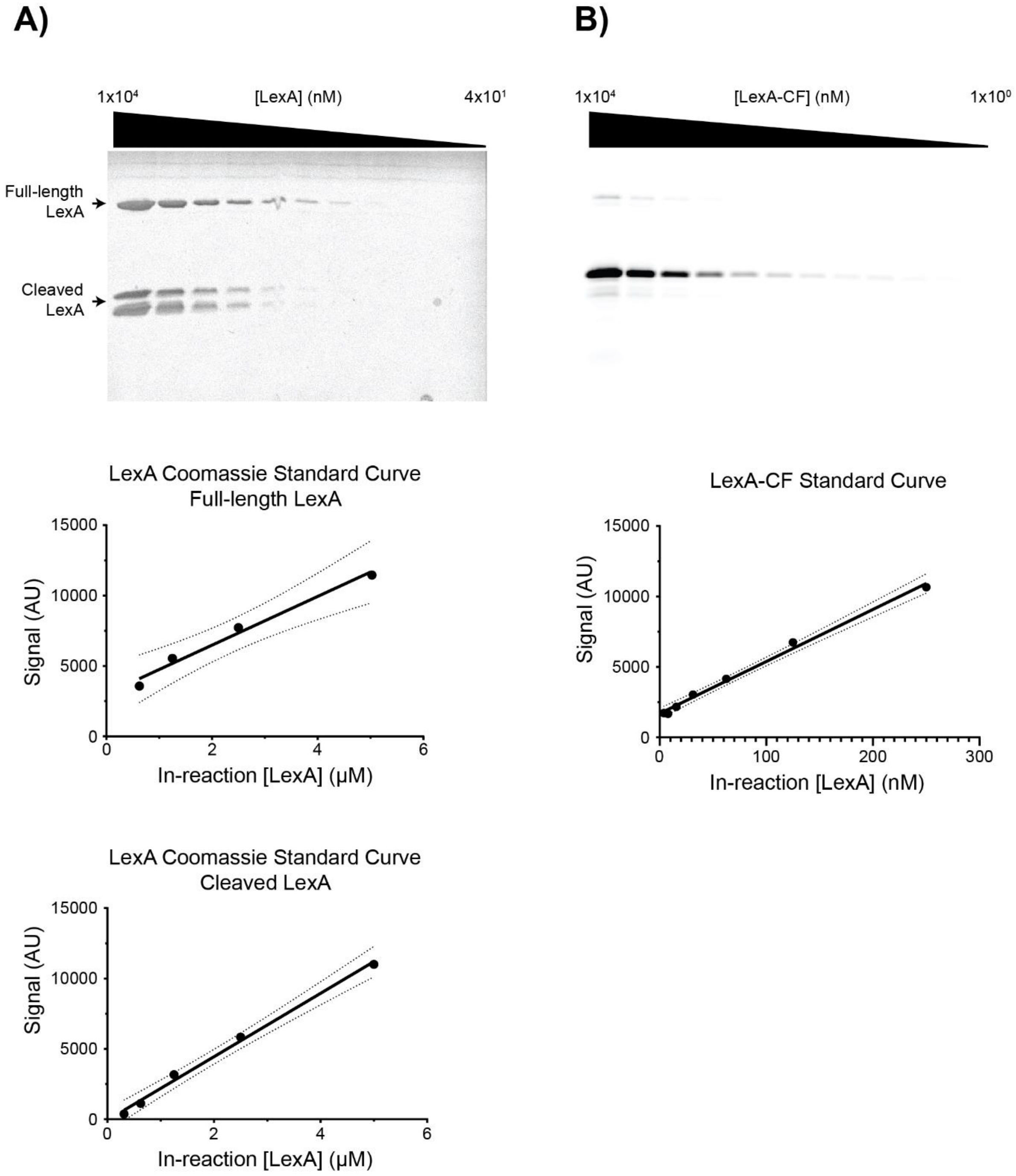


**Figure S2. Standard curves for quantitative cleavage assays. A)** LexA WT was auto-digested in alkaline autoproteolysis buffer for 3 hours at 37 °C and mixed with equimolar inactive LexA S119A. Samples were then serially diluted and loaded onto a 15% SDS-PAGE gel and stained with Coomassie. The linear range was determined by assessing the $R^2$ variation of at least four consecutive points along the curve as quantified after background correction by ImageJ. **B)** LexA-CF was auto-digested for 3 hours at 37 °C and loaded onto a 15% SDS-PAGE gel and imaged for fluorescence. Linear range was determined in an identical manner to LexA WT.

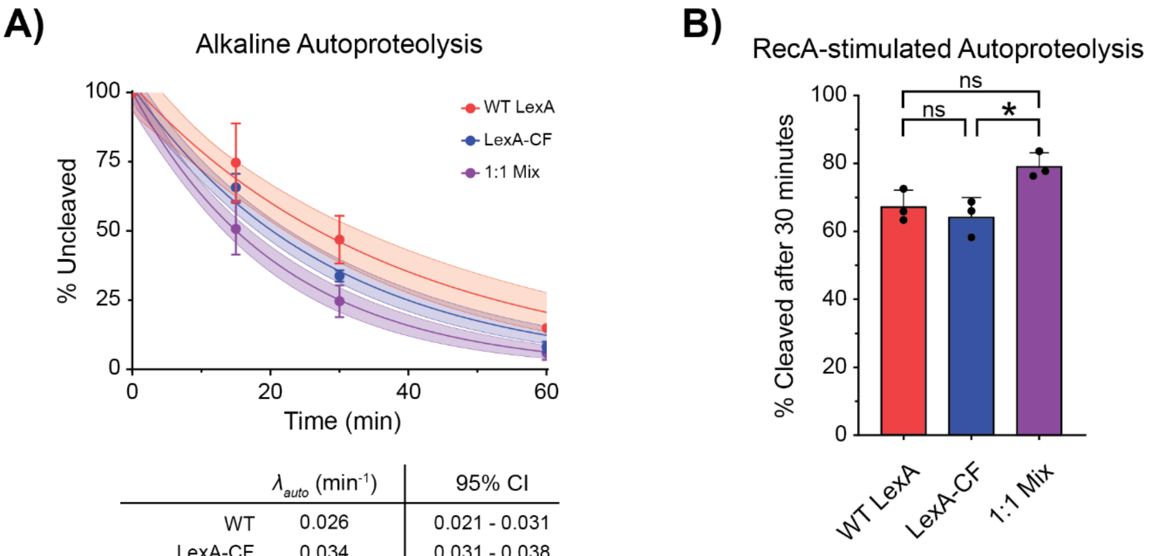


| | $\lambda_{auto}$ ($min^{-1}$) | 95% CI |
|---|---|---|
| WT | 0.026 | 0.021 - 0.031 |
| LexA-CF | 0.034 | 0.031 - 0.038 |
| 1:1 Mix | 0.046 | 0.041 - 0.052 |

**Figure S3. Label on LexA-CF does not affect autoproteolysis. A)** LexA WT, LexA-CF, and a 1:1 mix of both together were auto-digested in alkaline auto-proteolysis buffer for 60 minutes at 25 °C. Data was fit in GraphPad prism to a single-exponential decay (solid lines), with the points representing means and error bars representing standard deviations from three replicates. Shaded region shows the 95% confidence interval of the fit. Table shows the best fit and 95% confidence intervals for the rate of auto-proteolysis from the nonlinear fit. **B)** LexA-WT, LexA-CF, and a 1:1 mix of both together were mixed with pre-activated RecA WT (RecA*) and reacted for 30 minutes at 25 °C. Each data point from three replicates is shown with error bars representing the standard deviation of each. Data were compared using a one-way ANOVA assuming non-uniform standard deviations. The asterisk denotes a statistical significance of $p < 0.05$.

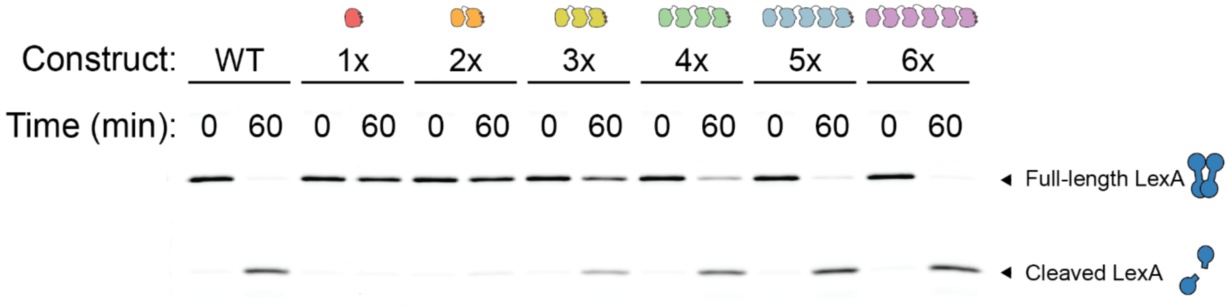


**Figure S4. Quantitative cleavage of LexA-CF.** Representative gel showing cleavage of 100 nM LexA-CF by each RecA construct after 60 minutes at 25 °C. Data was collected in triplicate and band intensities were quantified in ImageJ after background correction. The associated quantified data are summarized in **Figure 2B**.

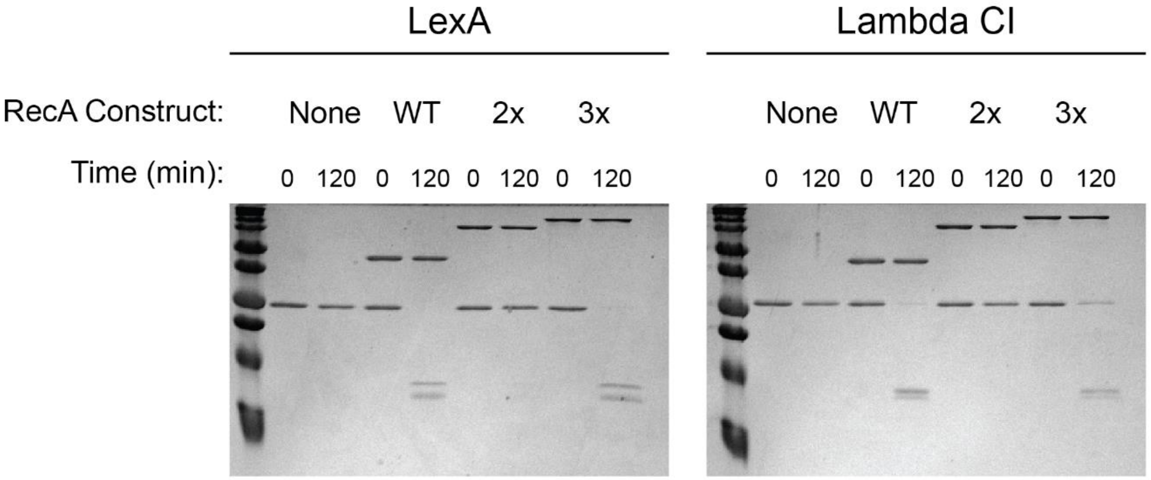


**Figure S5. RecA-stimulated auto-proteolysis of either LexA WT or λ phage repressor (CI).** Pre-activated RecA-WT (RecA*) was mixed with either LexA or CI and reacted for 120 minutes at 25 °C. Samples were then run on a 15% SDS-PAGE gel to visualize cleavage patterns.

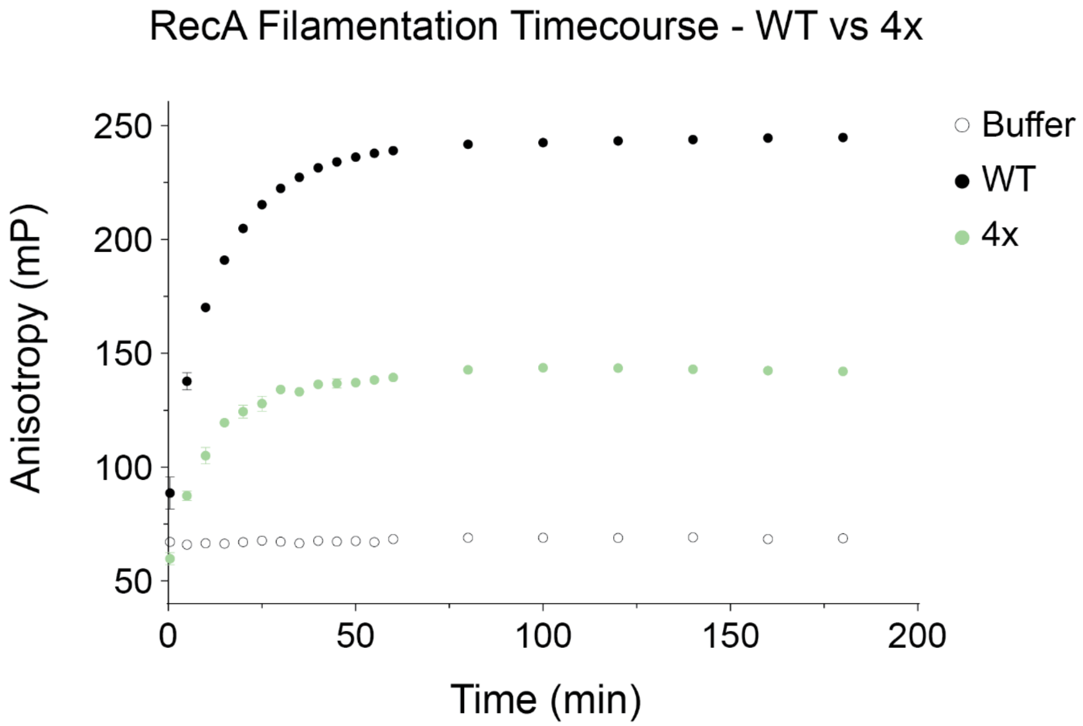


**Figure S6. Filamentation time course.** RecA WT, RecA4x, or no RecA (buffer control) were mixed with 2 nM 3'-FAM-labelled $(GGT)_6$ oligonucleotide under activation conditions and the anisotropy was monitored over time. Points are the mean from three replicates, with standard deviations that are too small for visible error bars.

**A)**

| Construct | Filamentation $K_D$ (nM) |
|---|---|
| WT; nucl. | 9600 (7200 - 13200) |
| WT; ext. | 130 (120 - 140) |
| 2x | — |
| 3x | 330 (290 - 380) |
| 4x | 70 (58 - 73) |
| 5x | 30 (25 - 36) |
| 6x | 15 (12 - 19) |

best-fit values from fit in terms of [construct]

With qualitative reaction conditions of:

$(GGT)_6$ = 1 μM

$[RecA]_{construct}$ = known; e.g. 250 nM for RecA4x

⇒

Estimate fraction bound to determine $[RecA^*]_{construct}$

| $[RecA^*]_{construct}$ (nM) |
|---|
| 180†† |
| — |
| 230 |
| 230 |
| 190 |
| 160 |

**B)**

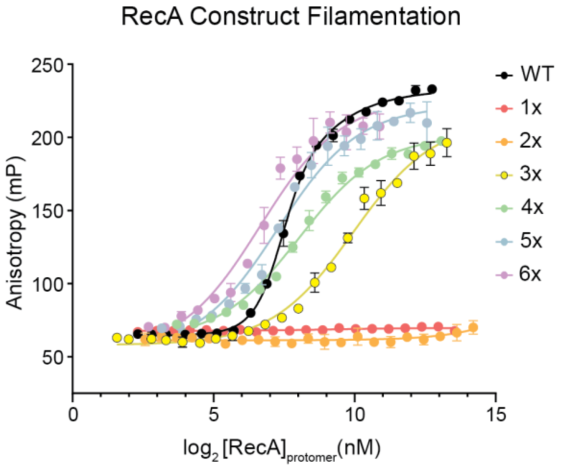


**C)**

| Construct | Filamentation $K_D$ (nM) |
|---|---|
| WT; nucl. | 9600 (7200 - 13200) |
| WT; ext. | 130 (120 - 140) |
| 2x | — |
| 3x | 990 (870 - 1100) |
| 4x | 260 (230-290) |
| 5x | 150 (130-190) |
| 6x | 100 (80-130) |

best-fit values from fit in terms of [total protomer]

With qualitative reaction conditions of:

$(GGT)_6$ = 1 μM

$[RecA]_{protomer}$ = known; 1 μM for all constructs

⇒

Estimate fraction bound to determine $[RecA^*]_{protomer}$

| $[RecA^*]_{protomer}$ (nM) |
|---|
| 810† |
| — |
| 380 |
| 600 |
| 680 |
| 730 |

Convert $[RecA^*]_{protomer}$ to $[RecA^*]_{construct}$ by:

$$[RecA_N^*]_{construct} = \frac{[RecA_N^*]_{protomer}}{N}$$

⇒

| $[RecA^*]_{construct}$(nM) |
|---|
| 180†† |
| — |
| 130 |
| 150 |
| 140 |
| 120 |

RecA WT concentrations calculated based on either the number of active protomers present in filaments† or the sum of filaments†† between two and six protomers in length (assuming a max of 6-unit filaments)

**Figure S7. Extended analysis of RecA filamentation. A)** RecA filamentation data from **Fig. 3A** were used to estimate the concentration of active RecA* in the quantitative LexA cleavage experiments. **B)** The data from **Fig. 3A** was also re-plotted based on the in-reaction concentration of RecA protomers, rather than $[RecA]_{construct}$, and re-fit. Data points are the means from three replicates with error bars showing standard deviation. Fits were performed using a non-linear fit to either quadratic binding or Adair model of cooperative binding (solid lines). **C)** The best-fit and 95% confidence intervals for $K_D$ are shown in the second column of the table. The best-fit $K_D$ values were then used to estimate the concentrations of active protomer and active construct in solution with the given transformations shown between columns of the table.

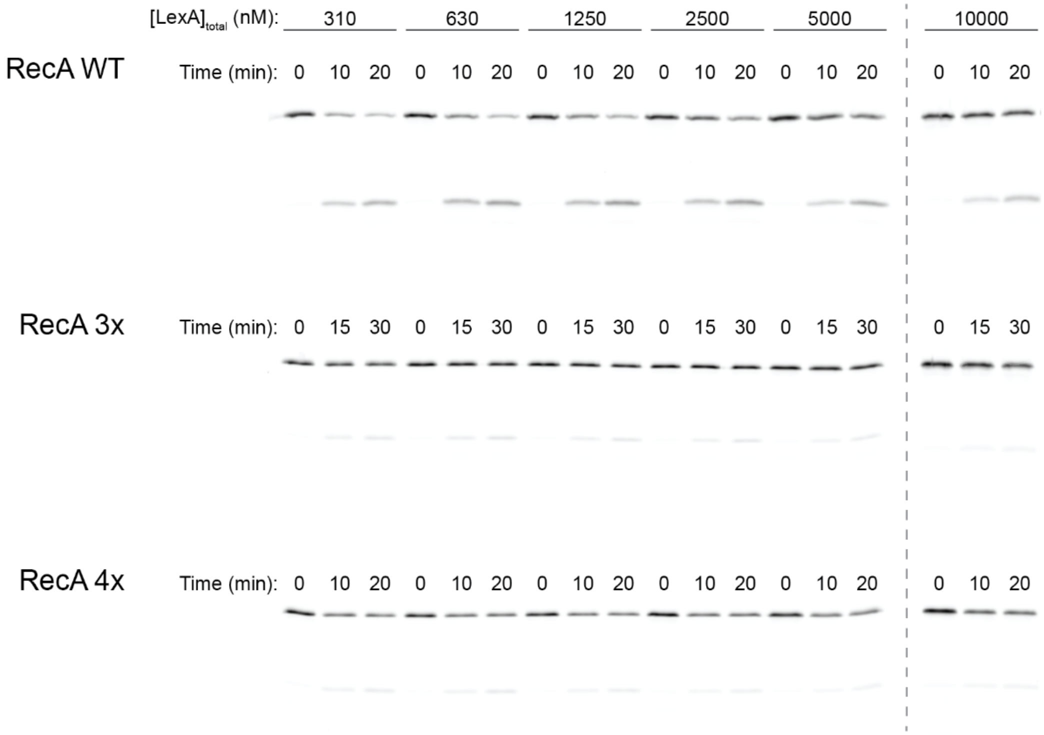


**Figure S8. Representative steady-state LexA cleavage gels.** Each gel is a representative replicate of RecA-WT, RecA3x, and RecA4x steady-state cleavage of LexA. Protomer concentrations in this assay were 600 nM for RecA3x and 450 nM for both RecA WT and RecA4x, with equimolar $(GGT)_6$. Each reaction contained a fixed amount of 100 nM LexA-CF with an increasing amount of unlabeled LexA-WT to yield the $[LexA]_{total}$ noted above each gel lane.

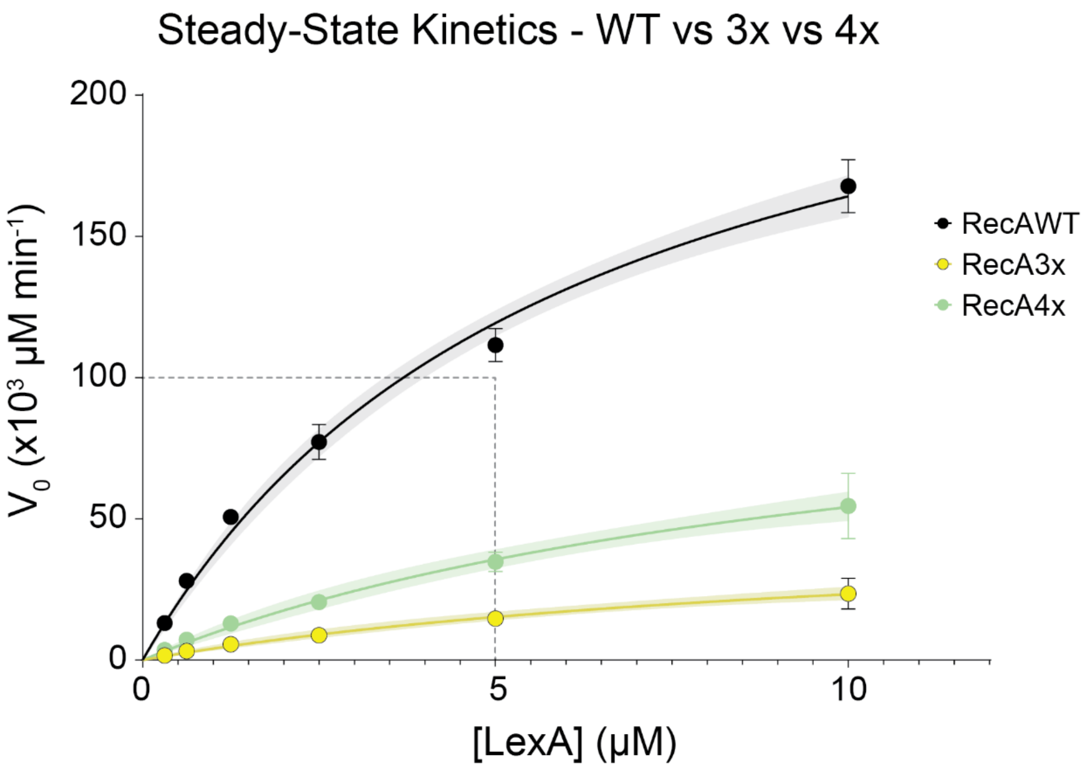


**Figure S9. Full steady-state kinetic curves for RecA constructs**. This graph shows the full quantified steady-state kinetic data for RecA-WT, RecA3x, and RecA4x. Points are the mean from three replicates and error bars are the standard deviation. Dotted lines show the region highlighted in **Figure 3C**. Shaded regions show the 95% confidence interval of each best-fit line shown in solid.

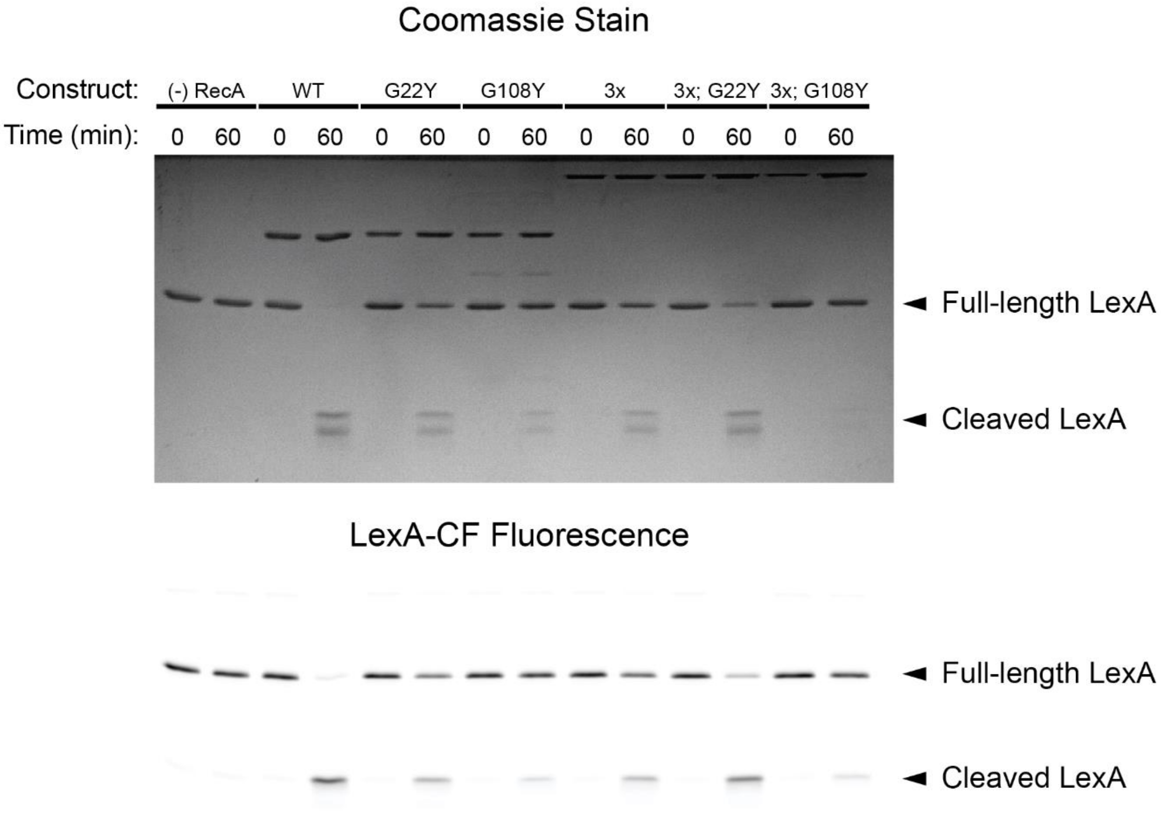


**Figure S10. LexA cleavage assessment for RecA mutants.** LexA mixture containing 1.9 μM unlabeled LexA and 100 nM LexA-CF was incubated with activated RecA constructs or no RecA. Samples were then loaded onto a 15% SDS-PAGE gel and imaged first for fluorescence (bottom panel) followed by Coomassie staining and reimaging (top panel).
